# A Developmentally Prometastatic Niche to Hepatoblastoma in Neonatal Liver mediated by the Cxcl1/Cxcr2 Axis

**DOI:** 10.1101/2021.10.22.465518

**Authors:** Li Fan, Qingfei Pan, Wentao Yang, Selene C. Koo, Cheng Tian, Liyuan Li, Meifen Lu, Anthony Brown, Bensheng Ju, John Easton, Sarangarajan Ranganathan, Soona Shin, Alexander Bondoc, Jun J. Yang, Jiyang Yu, Liqin Zhu

## Abstract

**Background and Rationale:** Hepatoblastoma (HB) is the most common pediatric liver cancer. Its predominant occurrence in very young children led us to investigating whether the neonatal liver provides a protumorigenic niche to HB development.

**Methods:** HB development was compared between orthotopic transplantation models established in postnatal day 5 and 60 mice (P5^Tx^ and P60^Tx^ models). Single-cell RNA-sequencing was performed using tumor and liver tissues from both models and the top candidate cell types and genes identified are investigated for their roles in HB cell growth, migration, and survival.

**Results:** We found that various HB cell lines including HepG2 cells were consistently and considerably more tumorigenic and metastatic in the P5^Tx^ model than in the P60^Tx^ models. Sc-RNAseq of the P5^Tx^ and P60^Tx^ HepG2 models revealed that the P5^Tx^ tumor was more hypoxic and had a larger number of activated hepatic stellate cells (aHSCs) in the tumor-surrounding liver which express significantly higher levels of *Cxcl1* than those from the P60^Tx^ model. We found these differences were developmentally present in normal P5 and P60 liver. We showed that the Cxcl1/Cxcr2 axis mediated HB cell migration and was critical to HB cell survival under hypoxia. Treating HepG2 P60^Tx^ model with recombinant CXCL1 protein induced intrahepatic and pulmonary metastasis and CXCR2 knockout in HepG2 cells abolished their metastatic potential in the P5^Tx^ model. Lastly, we showed that in metastatic HB patient tumors there was a similar larger population of aHSCs in the tumor-surrounding liver than in localized tumors, and tumor hypoxia was uniquely associated with HB patient prognosis among pediatric cancers.

**Conclusion:** We demonstrated that the neonatal liver provides a prometastatic niche to HB development via the Cxcl1/Cxcr2 axis.

## INTRODUCTION

Hepatoblastoma (HB) is a rare hepatic tumor of early childhood with unclear etiology.^1^ Accounting for only 0.5-2% of all cancer cases in children, however, the incidence of HB is rising globally at a pace faster than any other pediatric cancers.^2^ Although HB patients have an excellent five-year survival rate above 80%, children with advanced forms of HB, including those that have metastasized or relapsed, do not respond well to standard treatments and their five-year survival is less than 40%.^3^ There is clearly an urgent need to understand the biology behind the deadly forms of HB.

HB is one of the most genetically simple cancer types with mutational burdens as much as 10^4^-fold lower than those of adult cancers.^4, 5^ More than 80% of HB tumors only contain somatically acquired *CTNNB1* gene mutations,^6^ and HB initiation has been successfully modeled in genetic mouse models by activating *Ctnnb1* mutations in the embryonic liver.^7^ However, studies have repeatedly found that *Ctnnb1* mutations that are able to induce HB in the immature liver are non-oncogenic when activated in adult mouse liver,^8, 9^ suggesting that the early developing liver may provide a protumorigenic tumor microenvironment (TME) to HB development. Indeed, HB has a unique, tight association with early development even among solid tumors that affect only children – HB occurs almost exclusively in young children under the age of three. Very-low-birth-weight (VLBW) infants have a 20-50-fold higher risk of developing HB than those born with normal weight, and HB with unfavorable prognosis typically develops in children who are extremely premature at birth.^10, 11^ Based on these observations, we hypothesize that the immature liver provides a developmentally protumorigenic niche to HB development. Patient-derived xenografts (PDXs) and cell line-derived xenografts have been established for HB in multiple laboratories.^12, 13^ However, these models used adult mice as the host, and the HB tumors developed in these models rarely progress into the advanced, metastatic stage. In this study, we established a novel HB orthotopic transplantation model using postnatal day 5 (P5) mice as the host and then compared to mice transplanted at P60 to dissect the potential cellular and molecular contributors to the differences we observed.

## RESULTS

### P5 mouse liver is more prometastatic to human and mouse HB cells than the P60 liver

To assess the impact of the early developing liver on HB progression, we developed an orthotopic liver transplantation procedure using P5 mouse pups as the host (P5^Tx^ model) (**Figure 1A**). Due to their sensitivity to anesthetic, the small size of the liver, low post-surgery survival and dam rejection, we achieved an approximately 50% survival rate in this model. Liver transplantation was also performed in P60 mice in parallel (P60^Tx^ model). Tumorigenicity of two HB cell lines was compared between P5^Tx^ and P60^Tx^ models: HepG2, a validated human HB cell line^14, 15^ and HBS1, a mouse HB cell line derived from a previously reported Notch-driven HB genetic mouse model.^16, 17^ NSG mice were used as the host strain for HepG2 and CD-1 NU/NU nude mice for HBS1. We found that HepG2, when orthotopically transplanted at 5×10^4^ cells/mouse, developed lung metastasis in 10 out of the 13 P5^Tx^ mice while all the P60^Tx^ mice (15/15) grew tumors only in the liver (median survival = 40 and 43 days, respectively, *P* value = 0.038) (**Figure 1, B-D**). To ensure that the tumor cell number/gram of body weight or tumors cell leakage into the blood circulation did not account for the extensive metastasis in the P5^Tx^ model, we performed orthotopic transplantation at 5×10^5^ cells/mouse or tail vein injection up to 3×10^6^ cells/mouse in P60 mice and found no lung metastasis in either conditions six weeks post injection (data not shown). For HBS1, we have previously reported that this mouse HB line was metastatic in an adult CD-1 NU/NU (nude) mouse orthotopic allograft model.^17^ In this study, we found that the P5^Tx^ nude model consistently developed more lung metastases than the P60^Tx^ model when orthotopically transplanted with 5×10^4^ HBS1 cells/mouse and examined six weeks post transplantation while tumors in the liver grew at a similar rate (**Figure 1E, F**). HBS1 cells had a genetically engineered ZsGreen (ZsG) reporter gene. We noticed an evident presence of ZsG^+^ tumor cells in both near- and far-tumor liver (defined as <500 μm and >5mm away from the tumor border, respectively) in the P5^Tx^ mice but not in the P60^Tx^ mice (**Figure 1G**), suggesting a more successfully tumor cell dissemination in the former. We also tested HB214, a human HB cell line derived from a previously reported subcutaneous HB PDX model.^12^ HB214 was not tumorigenic when injected into P5 NSG mice at 5×10^4^ cells/mouse. When injected into P21 and P60 liver at 1×10^6^ cells/mouse, HB214 developed tumors in 5/5 P21^Tx^ mice with faithful HB histopathology while no tumors were found in P60^Tx^ mice (**Suppl Figure S1**). Although no lung metastasis developed in HB214 P21^Tx^ mice, this result supports the notion that the mouse liver at an earlier developmental stage is more protumorigenic than the adult liver.

**Figure 1.**
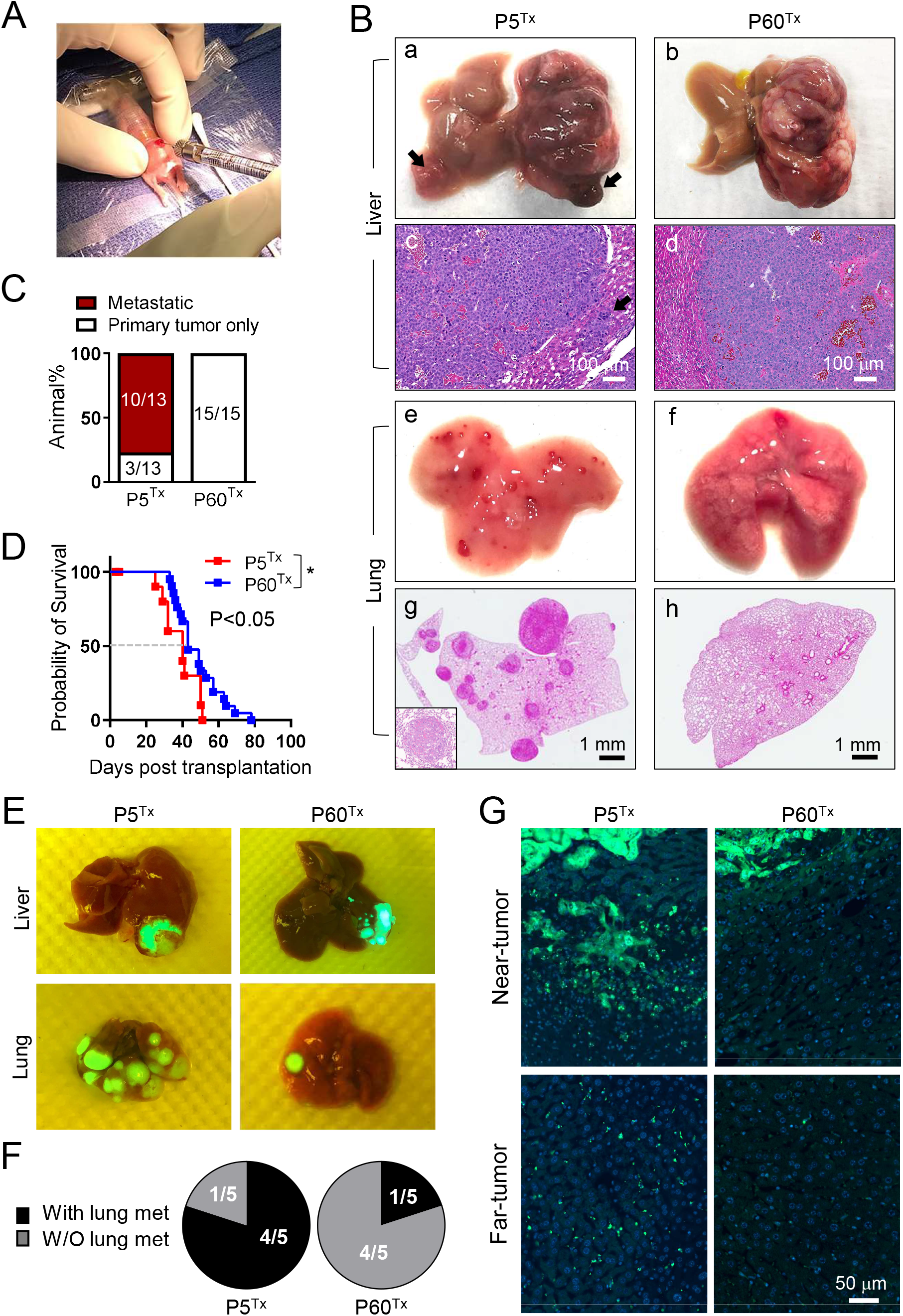
The neonatal liver promotes HB development. **(A)** Orthotopic liver transplantation using a P5 mouse pup. **(B)** The gross **(a, b, e, f)** and H&E **(c, d, g, h)** images of the liver and lung collected from the HepG2 P5^Tx^ and P60^Tx^ model. Arrows in **(a, c)**: intrahepatic metastases; inset in **(g)**: a higher magnification image of a lung metastasis. Scale bars as indicated. **(C)** Comparison of the lung metastasis rate between the HepG2 P5^Tx^ and P60^Tx^ model. **(D)** Kaplan-Meier animal survival curves of the HepG2 P5^Tx^ and P60^Tx^ model. **(E)** Merged gross/ZsG images of the liver and lung collected from the HBS1 P5^Tx^ and P60^Tx^ model. Liver tumors and lung metastases are ZsG^+^. **(F)** Pie chart comparison of the lung metastasis rate between the HBS1 P5^Tx^ and P60^Tx^ model. **(G)** ZsG fluorescence images of the near- and far-tumor liver in the HBS1 P5^Tx^ and P60^Tx^ model. All images share the same 50 μm scale bar.

### Single-cell RNA-seq reveals key differences in the tumor-surrounding liver between the P5^Tx^ and P60 ^Tx^ HepG2 models

To determine what contributed to the different metastasis outcome in the P5^Tx^ and P60^Tx^ models, we performed single-cell RNA-seq (sc-RNAseq) analysis of the tumor masses as well as tumor-surrounding liver tissues, or peritumoral liver, collected from a P5^Tx^ mouse and a P60^Tx^ mouse six weeks post HepG2 transplantation (**Figure 2A**). A total of 8881 HepG2 cells were readily identified from the P5^Tx^ and P60^Tx^ mice because of their human origin, and partitioned into 9 clusters without distinct segregation (**Suppl Figure S2A**). Sequencing quality of the HepG2 cells was confirmed by the high numbers of total genes and unique molecular identifiers (UMIs) detected, and the low number of mitochondrial genes (**Suppl Figure S2B**). Different from the largely homogeneous tumor cells, 14675 cells detected from the peritumoral liver tissues formed 17 well-segregated clusters (**Suppl Figure S2C**). Seven hepatic linages were annotated based on the scores of known lineage-specific gene signatures across the 17 clusters (**Figure 2B** & **2C**, **Suppl Figure S2D**, and **Suppl Table S1**).^18–20^ Expression of exemplar marker genes validated the lineage annotation by showing high cluster specificity for each lineage (**Suppl Figure 2E**). Sequencing quality of each hepatic lineage was similarly confirmed by the numbers of total genes, UMIs, and mitochondrial genes detected (**Suppl Figure 2F**).

**Figure 2.**
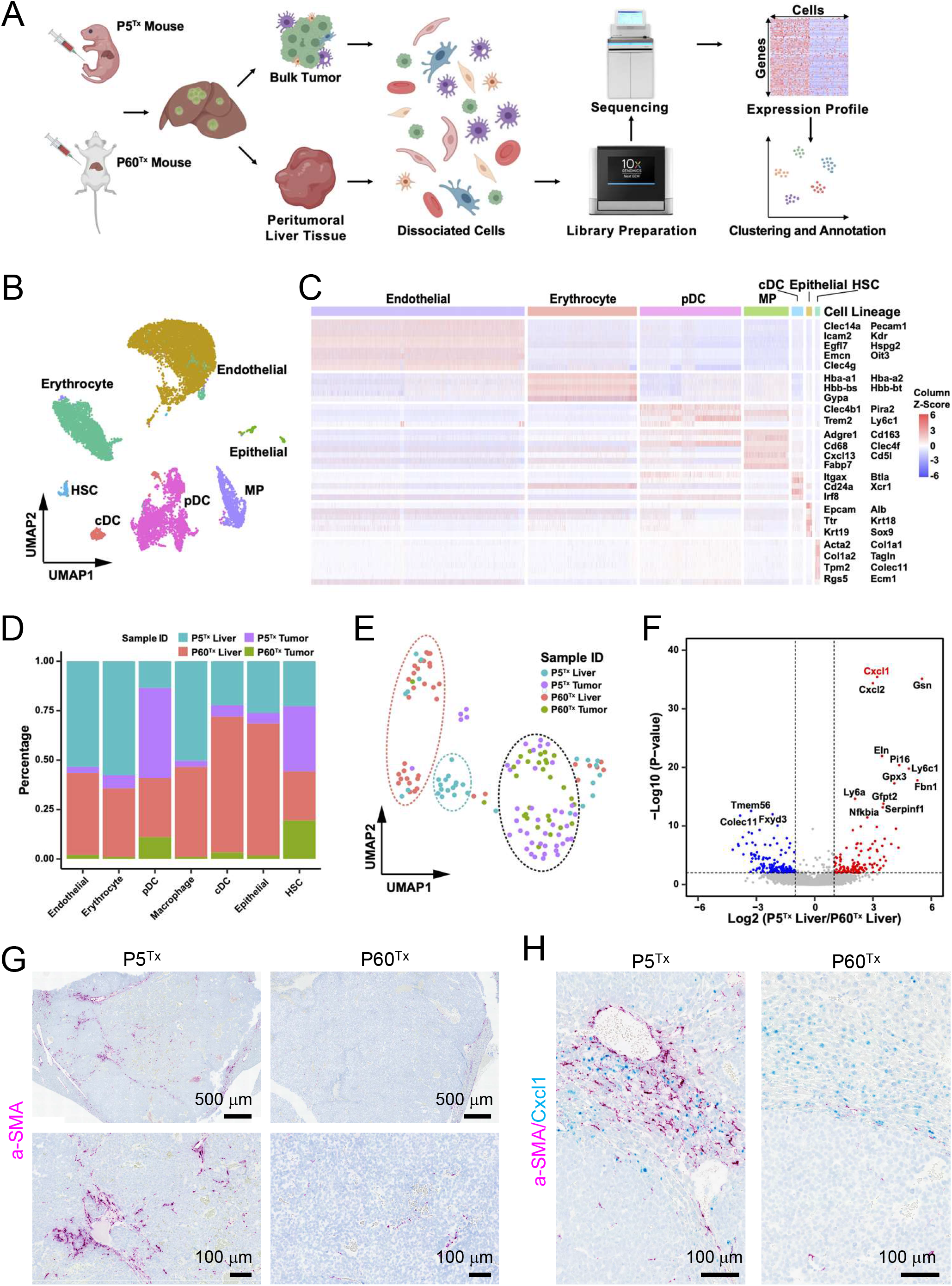
A scRNA-seq transcriptomics comparison of the tumor and liver tissues from the HepG2 models reveals a significant difference between the P5^Tx^ and P60^Tx^ peritumoral aHSCs. **(A)** The schematic illustrating the experimental design, single cell isolation, sequencing, and analysis. **(B)** Cell type annotation by the expression of marker gene signatures. HSC, hepatic stellate cell; cDC, conventional dendritic cell; pDC, plasmacytoid dendritic cell; MP, macrophage. **(C)** Heatmap showing the expression of marker gene signatures of each hepatic cell type. The columns denote cells, and the rows denote genes. **(D)** Bar plot showing the proportion of cells from the indicated sample sources across seven hepatic cell types. **(E)** UMAP illustration of the HSC clustering across the four indicated sample sources. The dashed ovals represent the main HSC clusters. (**F)** Volcano plot of differential expression analysis of the peritumoral HSCs from the P5^Tx^ and P60^Tx^ samples. Red dots indicate genes up-regulated in the P5^Tx^ liver HSCs and the blue dots indicate genes up-regulated in the P60^Tx^ liver HSCs. *P*-value <0.01 coupled with a Fold Change >2 or <0.5 were used to define the significantly differentially expressed genes. **(G)** RNAscope ISH of *αSMA* on the P5^Tx^ and P60^Tx^ tumors. Magenta: aSMA signal; blue: hematoxylin counterstain. Scale bars as indicated. **(H)** Dual RNAscope ISH of *αSMA/Cxcl1* on the P5^Tx^ and P60^Tx^ tumors. Magenta: *αSMA* signal; cyan: *Cxc11* signal, blue: hematoxylin counterstain. Scale bars as indicated.

Interestingly, hepatic stellate cells (HSCs), a well-known contributor to liver cancer metastasis,^21, 22^ showed a high level of tumor infiltration in both P5^Tx^ and P60^Tx^ models (**Figure 2D**). We found HSCs from the P5^Tx^- and P60^Tx^-peritumoral liver clustered separately while those found within the tumors were similar (**Figure 2E**). Among the top upregulated genes in the P5^Tx^ peritumoral HSCs compared to those in the P5^Tx^ model is a Cxcr2 ligands, *Cxcl1* (**Figure 2F**). Cxcl1/Cxcr2 axis has been shown to contribute to the metastasis of many adult solid tumors,^22–24^ but its role in pediatric solid tumors remains elusive. Another Cxcr2 ligand, Cxcl2, also showed higher expression in P5^Tx^ liver HSCs. To validate these findings, we performed dual *αSMA/Cxcl1* RNAscope in-situ hybridization (ISH) on tumor sections.

Cxcl1 is a small, secreted cytokine that is difficult to detect and localize via immunohistochemistry (IHC). Alpha-SMA is well recognized marker of activated HSCs (aHSCs).^25^ We confirmed that there was a larger number of αSMA^+^ aHSCs within the P5^Tx^ tumor (**Figure 2G**) as well as a larger population of *Cxcl1*-expressing aHSCs in the peritumoral liver than the P60^Tx^ model (**Figure 2H**). There were *Cxcl1*-expressing hepatocytes in both P5^Tx^ and P60^Tx^ models but *Cxcl1*-expressing aHSCs were predominantly found in the P5^Tx^ model. The heterogeneity of aHSC distribution and *Cxcl1* expression in- and outside the tumors made it challenging to perform quantitative comparisons between the two models. Nevertheless, our sc-RNAseq and histological analyses suggested that the aHSC population differed significantly between the P5^Tx^ and P60^Tx^ models.

### Activation and *Cxcl1* expression in HSCs differ significantly between the normal P5 and P60 liver

We suspected that the differences between aHSCs and their *Cxcl1* expression we detected in the P5^Tx^ and P60^Tx^ liver were part of developmental differences between the normal P5 and P60 liver. Therefore, we performed IHC and immunoblotting using HSC markers αSMA, desmin and vimentin on wildtype P5 and P60 (P5^WT^ and P60^WT^) NSG mouse liver. All three makers have been reported to increase during HSC activation.^26^ We found, indeed, there was a much larger population of αSMA^+^, desmin^+^ and vimentin^+^ cells as well as a higher expression of these proteins in the P5^WT^ liver than the P60^WT^ liver (**Figure 3A, B**). *Cxcl1* RNAscope ISH also showed more intense staining in the P5^WT^ liver than the P60^WT^ liver (**Figure 3C**). Dual RNAscope ISH of *αSMA/Cxcl1* showed that many *Cxcl1*-expressing cells in the P5^WT^ liver, although not all, colocalized with the *aSMA^+^* aHSCs (**Figure 3D**). To our knowledge, this is the first time that HSCs were shown to be markedly more active in the neonatal mouse liver than the adult, providing evidence that HB develops in a different TME than adult liver tumors.

**Figure 3.**
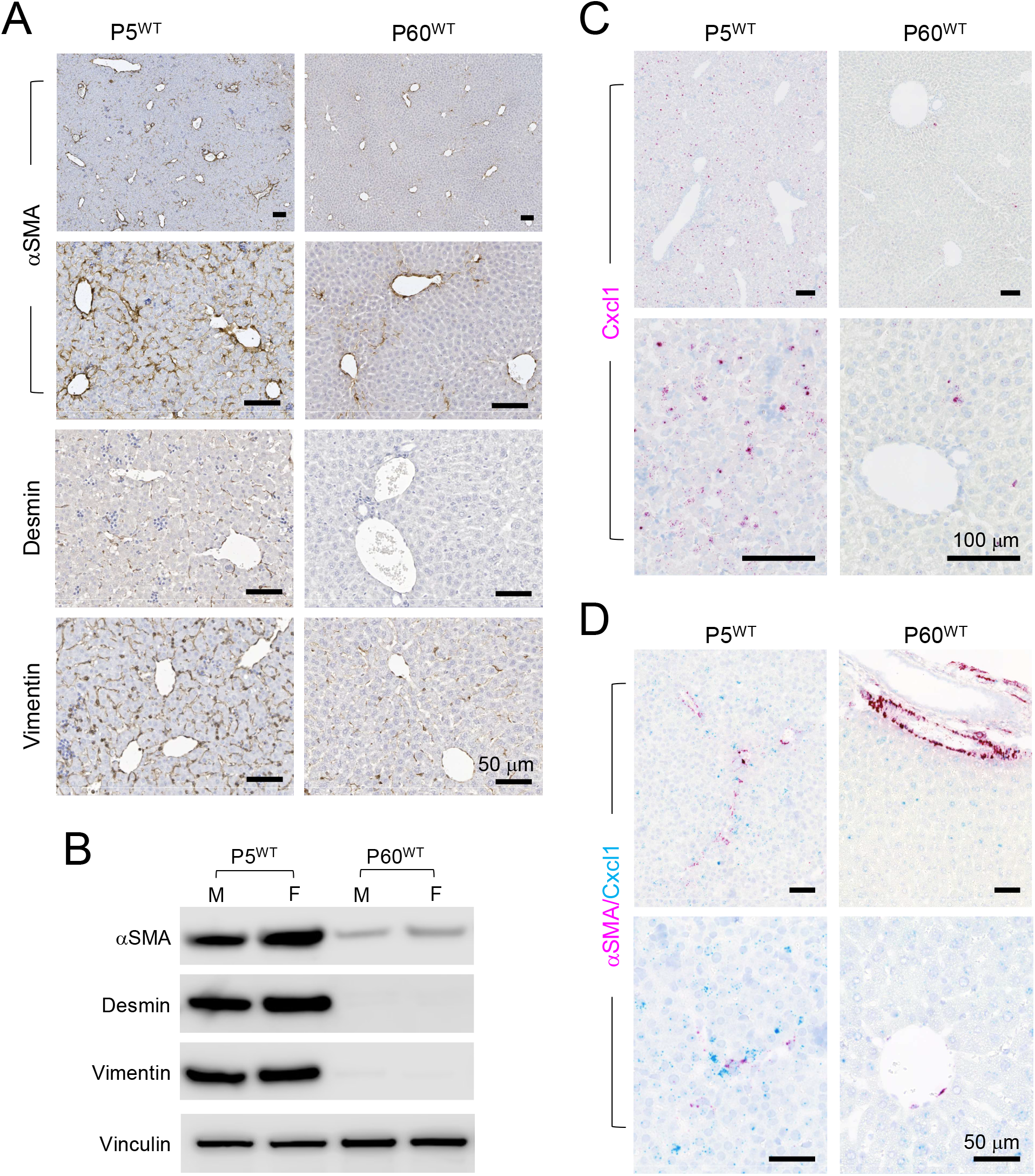
The wildtype P5 mouse liver has a much larger population of aHSCs than the P60 liver. **(A)** IHC of the indicated HSC markers on the P5^WT^ and P60^WT^ liver. All scale bars are 50 μm. **(B)** Immunoblots of the indicated HSC markers using whole cell proteins isolated from the P5^WT^ and P60^WT^ liver. **(C)** RNAscope ISH of *Cxcl1* on the P5^WT^ and P60^WT^ liver. Magenta: *Cxcl1* signal; blue: hematoxylin counterstain. All scale bars are 100 μm. **(D)** Dual RNAscope ISH of *αSMA/Cxcl1* on the P5^WT^ and P60^WT^ liver. Magenta: *αSMA* signal; cyan: *Cxc11* signal, blue: hematoxylin counterstain. All scale bars are 50 μm

### Peritumoral aHSCs promotes HB cell migration and dissemination in a CXCL1-dependent manner

Since there were no HSCs from neonatal human liver available, we utilized two human HSC lines – LX-2, an immortalized human HSC cell line, and primary human HHSteC cells (ScienCell) – both isolated from adult human liver to study their interaction with HB tumor cells. Although it is likely that neonatal HSCs are different from those in the adult liver beyond their activation status, we focused on answering the question whether peritumoral aHSCs would affect HB cell behaviors in a Cxcl1-dependent manner. Both HSC lines were propagated on plastic culture plates to maintain an activated state.^27, 28^ We then performed a cytokine array assay using conditioned medium (CM) from these HSCs as well as HepG2 and HB214 cells. We found that both HSC lines secreted significantly higher levels of CXCL1 than tumor cells (**Figure 4A**). ELISA using the CMs confirmed the higher secretion of CXCL1 by LX-2 and HHSteC (185 and 1290 pg/ml, respectively) than HepG2 and HB214 cells (2 and 5 pg/ml, respectively) (**Figure 4B**). We also found other cytokines that signaled through the same receptor CXCR2, including CXCL2/3, CXCL5, CXCL6 and CXCL8, were secreted at significantly higher levels in the HSCs than HB tumors cells (**Figure 4A and Suppl Figure S3**).

**Figure 4.**
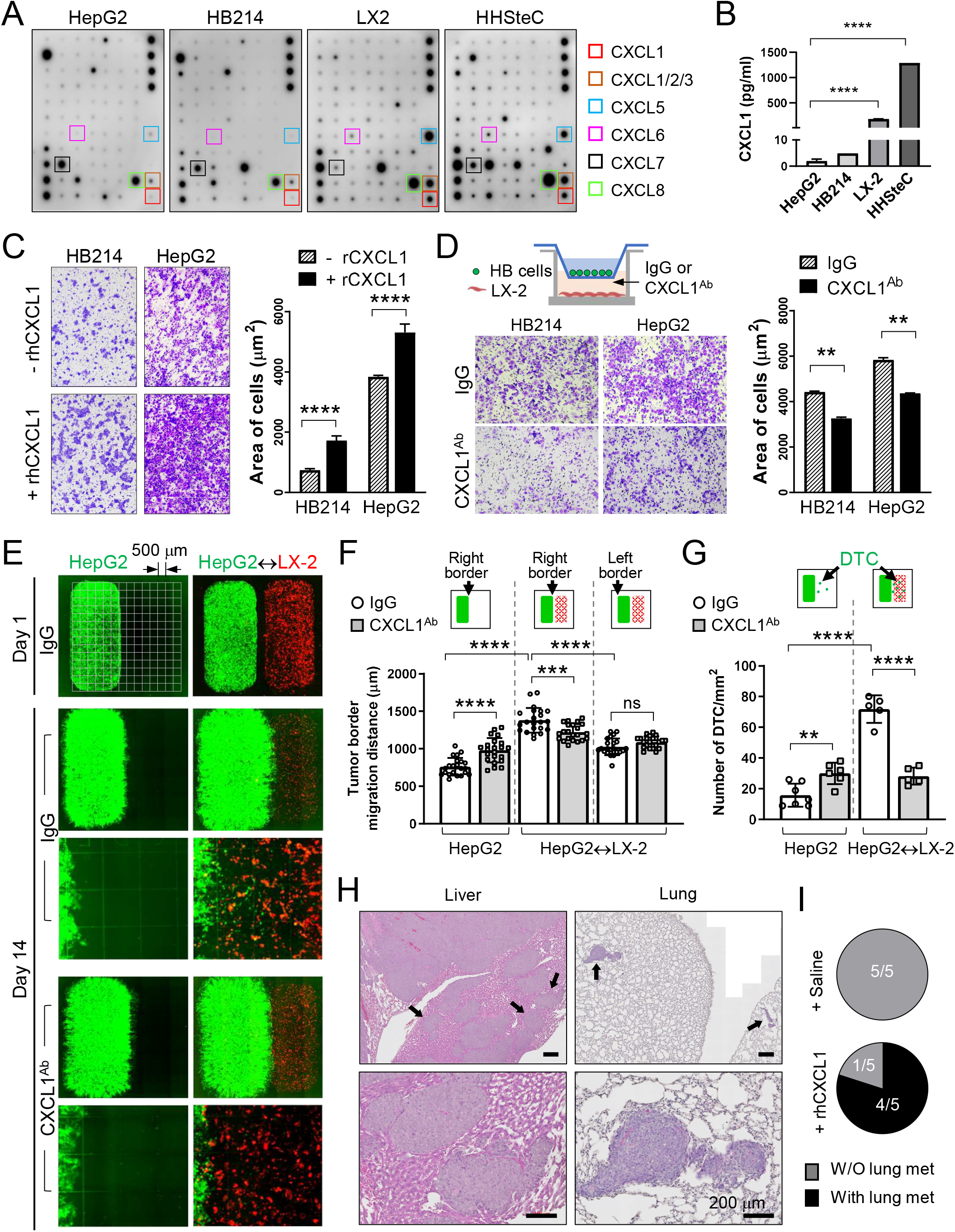
Peritumoral aHSCs promotes HB cell migration and dissemination in a CXCL1-dependent manner in vitro and in vivo. **(A)** Cytokine array assay using the CM from the indicated cells. Boxes: CXCR2 ligands. **(B)** CXCL1 Elisa using the CM from the indicated cells. **(C)** Images and quantification of the transwell migration assay of the HB214 and HepG2 cells pretreated with or without rhCXCL1. **(D)** Images and quantification of the transwell migration assay of the HB214 and HepG2 cells as indicated in the top diagram. LX-2 was cultured in the bottom chamber with IgG or CXCL1^Ab^ added in the medium. Number of cells/μm^2^ were counted and compared. **(E)** Day 1 and Day 14 GFP/RFP fluorescence images of the two-well cocultures of GFP^+^ HepG2 and RFP^+^ LX-2. HepG2 was placed in the left well with or without RFP^+^ LX-2 in the right chamber, and treated with IgG or CXCL1^Ab^. Culture slides with an imprinted 500-μm grid were used as indicated in the top left image. **(F)** Comparison of the HepG2 migration distance on the indicated border from Day 1 to Day 14 in **(E)** measured by using the 500-μm grid. **(G)** Comparison of the number of disseminated HepG2 cells/mm^2^ in the indicated conditions. **(H)** H&E images of the liver and lung of the P60^Tx^ model treated with rhCXCL1. Arrows: metastases I the liver and lung. All scale bars are 200 μm. **(I)** Pie charts showing the number of animals with or without lung metastasis in the saline- or rhCXCL1-treated P60^Tx^ model. All *p*-values were calculated by unpaired two-tailed *t*-test: ns, not significant, ** <0.01; *** <0.001; ****<0.0001.

When treated with recombinant human CXCL1 (rhCXCL1) protein, HepG2 and HB214 cells showed significantly increased migration ability in vitro (**Figure 4C**). Transwell migration assay with LX-2 placed in the bottom chamber also promoted the migration of both HepG2 and HB214 cells. Adding a neutralizing CXCL1 antibody (CXCL1^Ab^) in the bottom chamber suppressed this effect (**Figure 4D**). To better visualize peritumoral aHSC-induced HB cell migration, we set up a coculture system with LX-2 and HepG2 seeded separately by using a two-well silicone insert. Cell culture chambers with an imprinted 500-μm grid were used. GFP-expressing HepG2 cells were seeded in the left well (Well 1, or W1) with or without RFP-expressing LX-2 in the right well (W2) (**Figure 4E**). The insert was removed one day after seeding and cells were treated with IgG or CXCL1^Ab^. Tumor cell migration and dissemination were monitored and measured on Day 14. In IgG condition, HepG2 with LX-2 seeded in W2 showed significantly faster migration on the right border that faced LX-2, but not on the left border, than those seeded without LX-2 (average migration distance of the right border: 1381.0 μm vs 759.5 μm, respectively. *P* value < 0.0001) (**Figure 4E, F**). HepG2 with LX-2 in W2 also showed significantly greater cell dissemination from the right border than those without LX-2 (average number of disseminated cells: 16/mm^2^ vs. 72/mm^2^, respectively, *P* value < 0.0001). Upon CXCL1^Ab^ treatment, we observed a significant reduction in both the migration and dissemination of HepG2 cocultured with LX-2, indicating that LX-2-secreted CXCL1 functions as a chemoattractant to HepG2 (**Figure 4E, G**). Interestingly, we found HepG2 seeded without LX-2 showed a mild but significant increase in their migration and dissemination under CXCL1^Ab^ treatment, suggesting that the low level CXCL1 secreted by HB cells may also serve as a chemoattractant to keep tumor cells together.

To confirm the migration-promoting effect of CXCL1 on HB cells in vivo, we treated the P60^Tx^ model transplanted with 1×10^4^ /mouse HepG2 with weekly treatment of rhCXCL1 protein or saline. All mice were euthanized 60 days post transplantation. We found that 4 of the 5 mice treated with CXCL1 developed intrahepatic metastases and small but multifocal lung metastasis while no liver or lung metastasis was detected in the saline-treated mice (0/5) (**Figure 4H, I**). CXCL1^Ab^ treatment on the P5^Tx^ model was not performed due to their small size and low survival rate. Overall, these results showed that peritumoral aHSCs promotes HB cell migration and dissemination in a CXCL1-dependent manner.

### CXCR2 regulates the hypoxia response in HB tumor cells

Next, we tested the functional importance of the CXCL1 receptor, CXCR2, in HB development. We used CRISPR/Cas9 strategy to knockout (KO) the *CXCR2* gene in HepG2 and HB214 cells (**Figure 5A**). We found it difficult to obtain single-cell clones with complete *CXCR2* KO in both cell lines potentially because *CXCR2* was a one-exon gene and both cell lines had a low single-cell clonability. We selected one KO clone for each cell line that showed the most significant reduction in CXCR2 protein level and performed RNA-seq to determine the impact of *CXCR2* reduction on gene expression. Gene set enrichment analysis (GSEA) showed a significant downregulation of the hypoxia pathway in the *CXCR2^KO^* HepG2 cells (**Figure 5B**), a pathway that is well known for its prometastatic role in solid tumors.^29^ Indeed, the transcriptomics of HepG2 cells cultured under hypoxia showed a high degree of inversed correlation with that of the *CXCR2^KO^* HepG2 cells (**Figure 5C**). Possibly due to the low *CXCR2* expression in the parental HB214 cells (**Figure 5A**), *CXCR2^KO^* in HB214 caused a lesser degree of hypoxia-related gene expression changes which, however, still showed a similar inversed pattern compared to HepG2 cells cultured under hypoxia (**Figure 5D**), suggesting *CXCR2^KO^* in HB214 also negatively impacted hypoxia-responding genes. On the other hand, hypoxia did not induce consistent increases in *CXCL1* or *CXCR2* expression in the HB tumor cells and the two HSC cell lines cultured under hypoxia, indicating that these two genes are not direct downstream effectors of hypoxia (**Suppl. Figure S4**)

**Figure 5.**
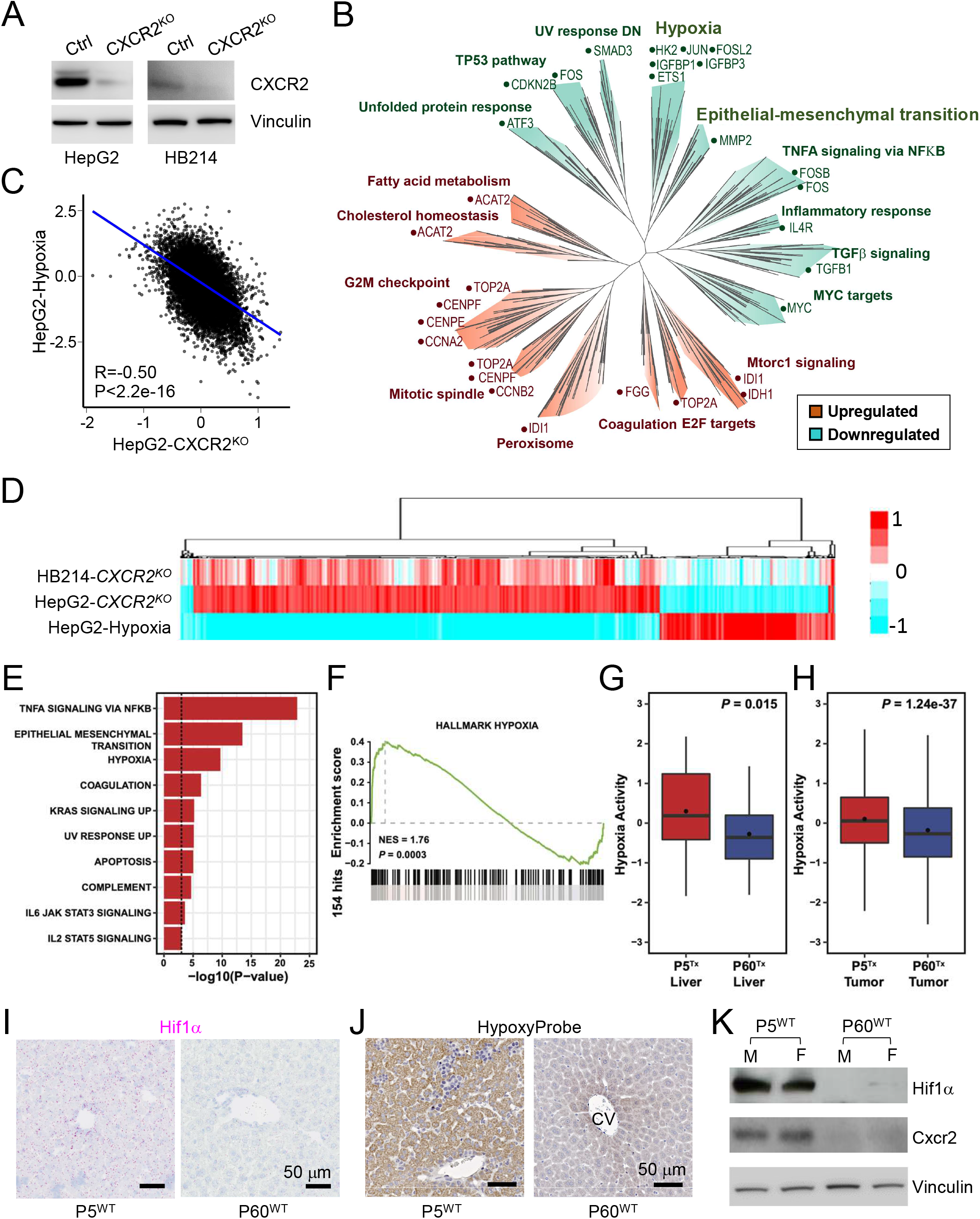
CXCR2 regulates the hypoxia response in HB tumor cells. **(A)** Immunoblots of CXCR2 and Vinculin in the indicated HB cells. **(B)** Pseudo-tree visualization of the GSEA comparing HepG2 control and *CXCR2^KO^* cells. **(C)** Spearman correlation of the RNAseq gene expression of the HepG2 cells cultured under hypoxia and *CXCR2^KO^* HepG2 cells. **(D)** Unsupervised clustering of the RNAseq gene expression of the indicated cell types. **(E)** Enrichment analysis of HALLMARK gene sets among the 140 up-regulated genes in P5^Tx^ liver HSCs using the Fisher’s exact test. Only those significantly enriched gene sets with *P*-value < 0.001 were shown. **(F)** GSEA of HALLMARK_HYPOXIA comparing the P5^Tx^ and P60^Tx^ liver HSCs. **(G, H)** Boxplot showing the distribution of hypoxia activities inferred by NetBID2 between the P5^Tx^ and P60^Tx^ peritumoral HSCs **(G)** and HepG2 tumor cells **(H)**. *P*-values were calculated by unpaired two-tailed *t*-test. **(I)** RNAscope ISH of *Hif1α* on the P5^WT^ and P60^WT^ mouse liver. Magenta: *Hif1α* signal; blue: hematoxylin counterstain. Both scale bars are 50 μm. **(J)** HypoxyProbe IHC on the P5^WT^ and P60^WT^ mouse liver. Both scale bars are 50 μm. **(K)** Immunoblots of Hif1α, Cxcr2, and vinculin using the whole proteins from the P5^WT^ and P60^WT^ mouse liver. M: male mouse; F: female mouse.

Based on these results, we looked into the scRNA-seq data of the HepG2 P5^Tx^ and P60^Tx^ models for hypoxia-related differences. Indeed, the hypoxia pathway is one of the top upregulated pathways in P5^Tx^ liver HSCs compared to those in P60^Tx^ liver (**Figure 5E-G**). HepG2 cells from the P5^Tx^ model also showed a mild but significant increase in hypoxia network activity than those from the P60^Tx^ model (**Figure 5H**). When examining the P5^WT^ and P60^WT^ mouse liver, we found the former showed a much higher expression of Hif1α – a master regulator of hypoxia^30^ – than the latter (**Figure 5I**). HypoxyProbe assay further confirmed higher hypoxia activity in P5^WT^ liver than P60^WT^ liver (**Figure 5J**). Immunoblotting confirmed higher protein levels of Hif1α and CXCR2 in P5^WT^ liver than P60^WT^ liver (**Figure 5K**). Overall, our data suggest that the neonatal liver is developmentally more hypoxic compared to adult liver, rendering a high hypoxia activity in its HSCs as well as tumor cells growing within.

### CXCL1/CXCR2 axis is essential to HB cell migration and survival under hypoxia

Hypoxia is known to be tightly associated with epithelial-mesenchymal transition (EMT). Indeed, HepG2 cells cultured under hypoxia showed a significant upregulation of the Hallmark_EMT pathway, which was downregulated in *CXCR2^KO^* HepG2 cells and HB214 cells although the latter did not reach statistical significance (**Figure 6A**). In vitro migration assay confirmed that HepG2 and HB214 cells cultured under hypoxia had a significant increase in their migration ability, and adding rhCXCL1 further increased migration ability of the hypoxic cells (**Figure 6B**). We noticed that hypoxia also had a strong, negative impact on the survival of HB cells. HepG2 and HB214 cultured under hypoxia for seven days showed a dramatic increase in their apoptosis which, however, was largely reversed by rhCXCL1 treatment (**Figure 6C**). Based on these observations, we hypothesized that CXCL1-secreting HSCs could protect HB cells from hypoxia-induced apoptosis, an effect which would be lost to *CXCR2^KO^* HB cells. To test this, we cultured GFP^+^ control and *CXCR2^KO^* HepG2 and HB214 cells with or without LX-2 HSCs under normoxia and hypoxia. Tumor cell survival was monitored via their GFP fluorescence. No difference was seen in tumor cell survival when these cocultures were maintained under normoxia for two weeks (**Figure 6D, e-l**). As expected, hypoxia induced a significant cell death in both the control and *CXCR2^KO^* HB cells grown without LX-2 after two weeks (**Figure 6D, m, q, o, s**). When cocultured with LX-2 under hypoxia, the control cells showed a significantly better survival (**Figure 6D, n, p**), while no improvement in cell survival was seen in the *CXCR2^KO^* HB cells (**Figure 6D, r, t**). To determine the importance of CXCR2 to HB development in vivo, we transplanted the control and *CXCR2^KO^* HepG2 cells in P5 mouse liver at 1× 10^4^/mouse and euthanized all animals after six weeks. We found *CXCR2^KO^* HepG2 cells generated significantly smaller tumors in the liver than the control cells and developed no lung metastasis (0/4), while 3/3 P5^Tx^ mice transplanted with the control HepG2 developed multifocal intrahepatic metastasis and 2/3 had lung metastases (**Figure 6E**). These results reveal the importance of CXCL1/CXCR2 axis-mediated HB-aHSCs interaction in HB cell migration and survival under hypoxia.

**Figure 6.**
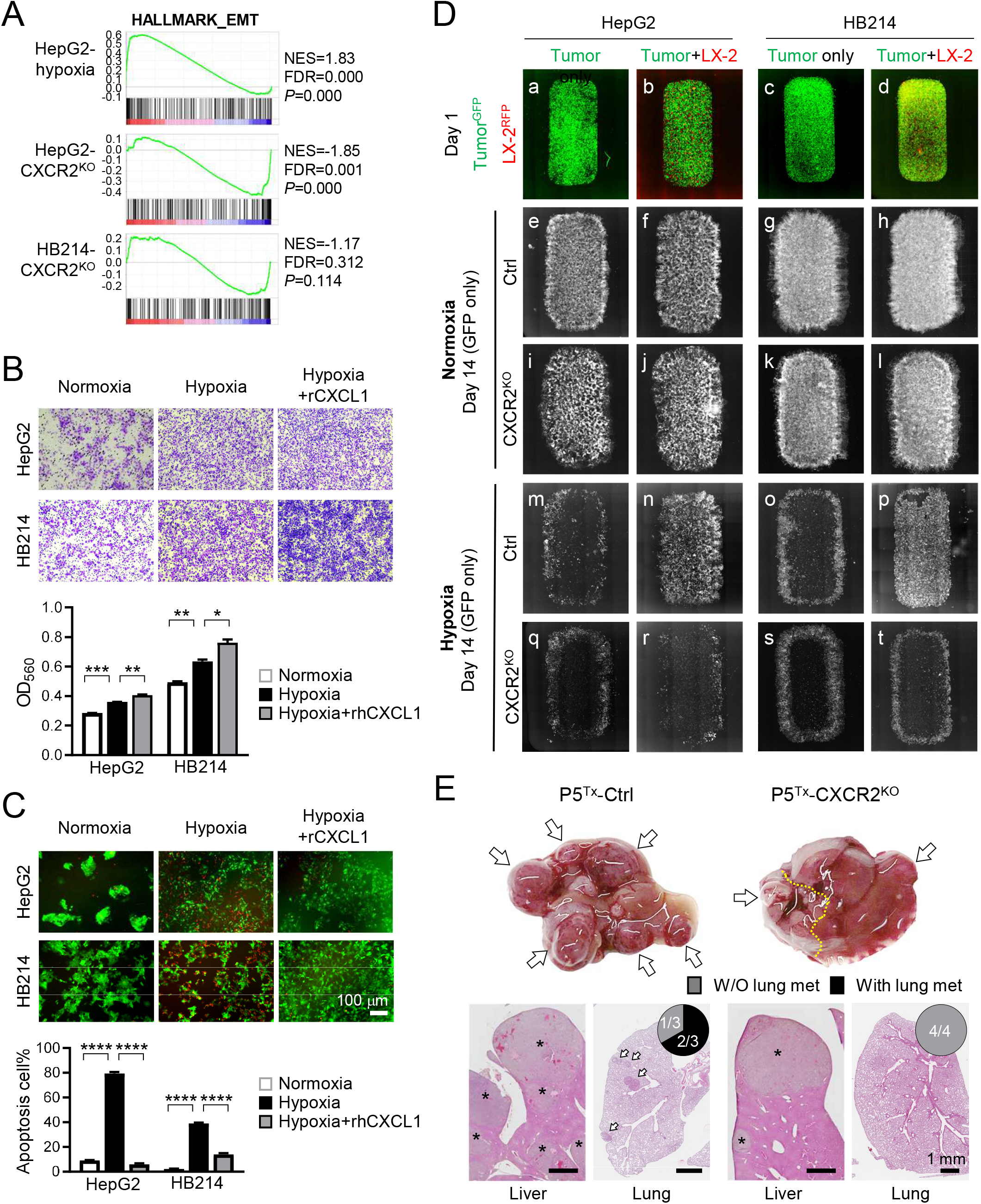
CXCL1/CXCR2 axis is essential to HB cell migration and survival under hypoxia. **(A)** GSEA of HALLMARK_EMT gene set by comparing the RNAseq profiles of the indicated cells with their control cells. **(B)** Transwell migration assay of the HepG2 and HB214 cells cultured under hypoxia. **(C)** Images and comparison of Caspase-3 apoptosis assay of the HepG2 and HB214 cells cultured under the indicated conditions. Apoptotic cells were labeled by NucView 530 RFP fluorescence. *P*-values were calculated by unpaired two-tailed *t*-test. **(D)** Day 1 and Day 14 fluorescence images of the GFP^+^ HepG2 and HB214 cocultured with or without RFP^+^ LX-2 under normoxia and hypoxia. Day 1 images are merged GFP/RFP and Day 14 images are GFP only. **(E)** Liver gross images (top) and H&E images of the liver and lung (bottom) of the P5^Tx^ model *CXCR2^KO^* transplanted with control or *CXCR2^KO^* HepG2 cells. * tumors in the liver; arrows: lung metastases. Pie charts indicate the incidence of lung metastasis. All scale bars are 1 mm.

### Peritumoral aHSCs and tumor hypoxia are associated with HB patient prognosis

To validate our aHSC-related findings in HB patients, a small group of Stage I (n=3) and Stage IV (n=3) HB patient tumors resected with >= 5mm of adjacent liver were identified. The tumor number was low due to the rareness of this pediatric cancer, and the fact that most of the HB surgical specimens in our archives were biopsies that did not have sufficient sampling of the tumor-liver interface. Because nearly all HB patients with metastatic disease receive chemotherapy prior to tumor resection, all three Stage IV tumors we examined in this study were post chemotherapy. To minimize the impact of HSC activation induced by chemotherapy in our comparison, the Stage I tumors chosen were also post treatment.

IHC for αSMA was performed to detect aHSCs in these patient tumors. We found that the Stage IV, metastatic HB tumors had consistently higher numbers of αSMA^+^ aHSCs in both the tumor core as well as the near-tumor liver than the Stage I, localized tumors (**Figure 7A**). Additionally, all three metastatic tumors showed more intense αSMA staining and more αSMA^+^ aHSCs in the near-tumor liver than in the tumor core, suggesting that peritumoral aHSCs may have a stronger association with HB metastasis than intratumoral HSCs. To validate our hypoxia-related findings in HB patients, we examined *HIF1A* expression in association with HB tumor risk using a published HB patient tumor transcriptomic database.^6^ We found *HIF1A* expression was significantly higher in the high-risk HB patient tumors than the intermediate- and low-risk ones (*P*=0.036 and 0.001, respectively) (**Figure 7B**). HB patients with lower *HIF1A* expression also had significantly longer survival (*P*=0.031) (**Figure 7C**). Since there have been multiple large transcriptomics studies on pediatric cancers, we compared the association between *HIF1A* expression in the tumor cells and patient survival in 12 different pediatric cancers including HB by pooling multiple publicly available RNAseq transcriptomics databases (see details in Methods). We found that HB was the only pediatric solid tumor that showed a significant association between tumor hypoxia activity and patient survival (Hazard ratio, or HR, = 10.12, *P*=0.015) (**Figure 7D**). Pediatric acute lymphoblastic leukemia was another pediatric cancer that showed significance, but with a much lower HR than HB (1.34 vs. 10.12). Compared to the other 33 adult cancer types we similarly calculated the association between *HIF1A* expression and patient survival, HB was still the only cancer type that had a HR above 2 (**Suppl Figure S5**). Taken together, these results indicate that hypoxia has a very unique, strong impact on HB development among pediatric cancers.

**Figure 7.**
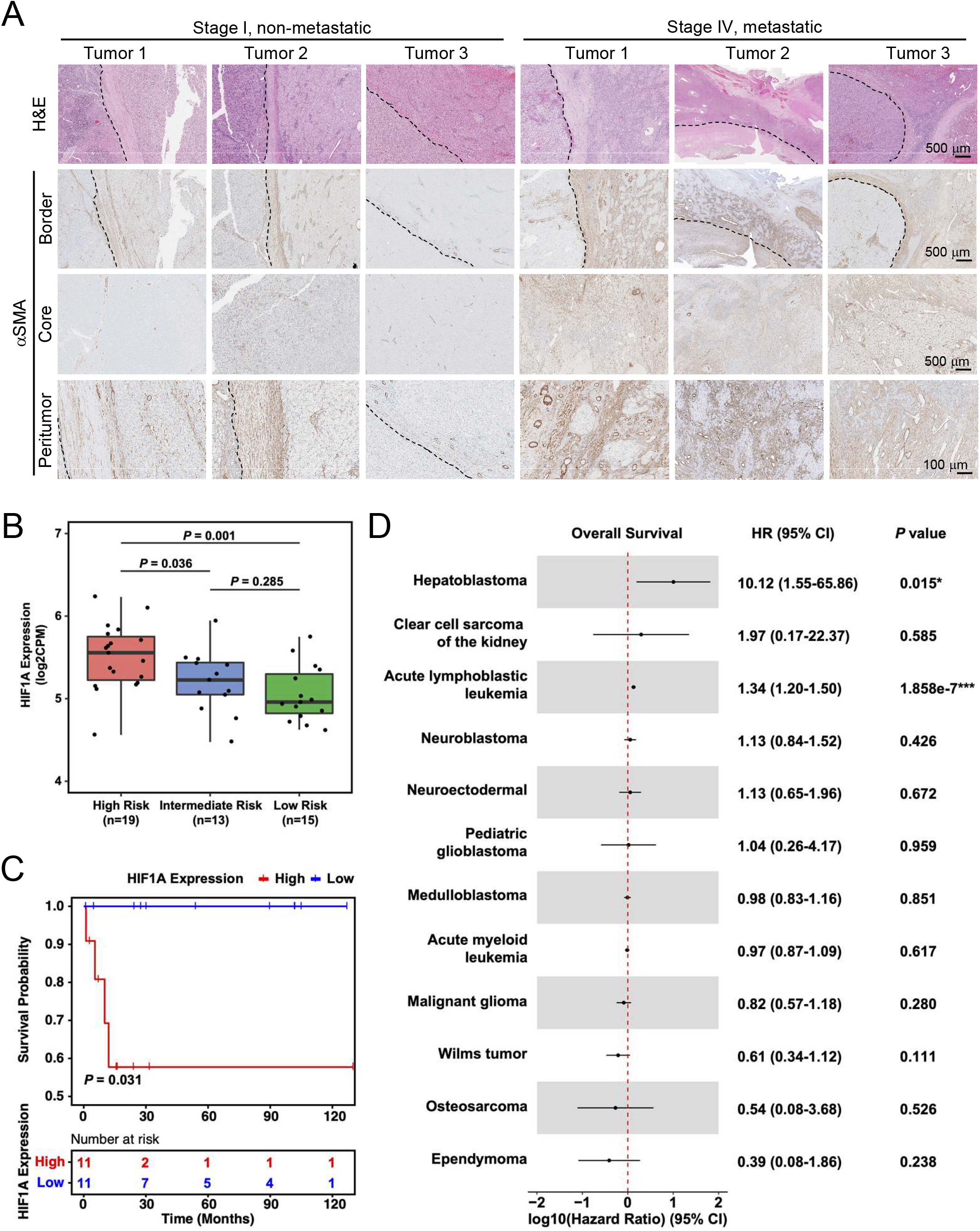
Peritumoral aHSCs and tumor hypoxia are associated with HB patient prognosis. **(A)** H&E and aSMA IHC images of HB patient tumors. Dashed lines: tumor border. Images on the same row share the same scale bar. **(B)** Boxplot showing the distribution of *HIF1A* expression among high-, medium-, and low-risk HB patient tumors. *P* values were calculated by unpaired two-tailed *t*-test. **(C)** Kaplan-Meier estimates of overall survival comparing the top (“High”) and bottom (“Low”) quarter of HB patients with regard to *HIF1A* expression. *P*-value was calculated by log-rank test. Tick marks indicate censoring. **(D)** Forest plot showing the association between *HIF1A* expression and overall survival among the patients of 12 types of pediatric cancer. HR, 95% CI and *P* values were calculated by univariate Cox proportional-hazards model.

## DISCUSSION

In this study, we compared HB metastasis outcome in mice transplanted with human and mouse HB cells at P60 and P5 and found that these cells were markedly more metastatic in the latter. Via single-cell transcriptomic analysis, we found higher levels of *Cxcl1* in the peritumoral aHSCs in the P5^Tx^ model than the P60^Tx^ model. We showed that this difference was part of the normal liver development that the P5^WT^ liver had a much larger population of aHSCs and higher expression of *Cxcl1* than the P60 ^WT^ liver. We showed that aHSCs secreted high levels of CXCL1, which was chemoattractant and prometastatic to HB cells both in vitro and in vivo. We found the P5^WT^ liver was indeed significantly more hypoxic than the P60^WT^ liver. Hypoxia induced HB cell migration but had a negative impact on survival, and aHSCs protected HB survival under hypoxia via the CXCL1/CXCR2 axis. Knockout of the CXCL1 receptor, *CXCR2*, in HB tumor cells compromised their hypoxia response and abolish their metastasis potential in the P5^Tx^ model. Using a limited number of patient tumors, we showed that there was a potential association between the number of peritumoral aHSCs and metastasis in HB patients. Lastly, by pooling transcriptomics database from 12 different types of pediatric cancers, we found, for the first time, that hypoxia has a unique, strong association with the poor survival of HB patients.

To our knowledge, work presented here is one of the first studies that places a pediatric solid tumor in its correct TME and demonstrates a strong impact of the developmentally immature TME on its progression. In fact, HB is the only pediatric cancer with a significantly increased risk in children and young adults born preterm.^31^ One common problem in premature infants is intermittent hypoxia in multiple organs. ^32, 33^ However, the pathophysiologic impact of intermittent hypoxia has been mainly studied in the respiratory and nervous systems with little knowledge of its impact on early liver biology and pathogenesis. Our results on CXCL1/CXCR2 and hypoxia provide a potential explanation for the strong association between HB risk and premature birth.^10^ Indeed, our transcriptomic analysis on 12 types of pediatric cancers revealed that HB is the only cancer type in which *HIF1A* expression is strongly associated with poor prognosis. We acknowledge that tumor metastasis we focused on in this study is only one of the contributing factors to HB prognosis. Further investigations on how hypoxia impacts liver and HB biology are in order to better understand HB development in young children.

We acknowledge that a major limitation of this study is the use of immunocompromised mice, which excluded the contribution from immune-related players that utilize the CXCL1/CXCR2 axis.^34–36^ We are bound by the limited research resources for HB.^37^ HepG2 cells are the only commercially available cell line for HB, and they are tumorigenic only in NSG mice. For our mouse HBS1 cells, our attempt to establish allograft model in immunocompetent B6 pups was unsuccessful due to their extremely poor post-surgery survival. A thorough investigation of the CXCL1/CXCR2 axis in various cellular compartments in HB patient tumors and tumor-surrounding liver will be necessary in the future to fully appreciate the involvement of this signaling pathway in HB development. We also acknowledge the small number of patient tumors we included due to the rarity of this cancer and the extremely limited specimens that have tumor-surrounding liver attached, which has made it difficult to establish statistical significance for our observations.

Rare pediatric cancers present one of the greatest challenges to the oncology community. Although most pediatric cancers have an excellent prognosis, their advanced forms still kill. With ever increasing appreciation of the importance of TME in adult cancers, it is necessary to begin the effect to investigate the developmental TME in pediatric cancers in order to achieve a more comprehensive understanding of their biology and more effective treatment for advanced diseases.

## METHODS

See supplemental materials.

## Acknowledgement

We thank Dr. Stefano Cairo (XenTech) for providing HB214 cell line.

## ABBREVIATIONS

HB: hepatoblastoma
HSC: hepatic stellate cells
aHSC: activated HSCs
TME: tumor microenvironment
VLBW: very-low-birth-weight
sc-RNAseq: single-cell RNA-sequencing
PDX: patient-derived xenograft
ZsG: ZsGreen
GFP: green fluorescent protein
IHC: Immunohistochemistry
ISH: in-situ hybridization
CM: conditioned media
rhCXCL1: recombinant human CXCL1
KO: knockout
GSEA: gene set enrichment analysis
EMT: epithelial-Mesenchymal Transition
ALL: acute lymphoblastic leukemia
RFP: red fluorescent protein

