## Supplementary methods, figures and tables for "A Developmentally Prometastatic Niche to Hepatoblastoma in Neonatal Liver mediated by the Cxcl1/Cxcr2 Axis"

**SUPPLEMENTARY MATERIAL**

**METHODS**

**Animals**

Animal protocols were approved by the St. Jude Animal Care and Use Committee. All mice were maintained in the Animal Resource Center at St. Jude Children's Research Hospital (St. Jude). Mice were housed in ventilated, temperature- and humidity-controlled cages under a12-hr light/12-hr dark cycle and given a standard diet and water *ad libitum*.

1. **$P5^{Tx}$ and $P60^{Tx}$ HB orthotopic transplantation models**
   HepG2 cells were surgically injected into the liver of P5 and P60 male and female NSG mice (JAX). Briefly, Mice were anesthetized with 2-3% isoflurane/O2, positioned ventrally, and secured on a heating pad. A midline longitudinal abdominal incision was made to expose liver left lobe and HepG2 cells were injected at $5\times10^4$/mouse in 2 µl cold growth factor-reduced (GFR) matrigel (Corning) using a 5-µl Hamilton syringe and a 27-gauge needle (Hamilton Company, Bonaduz, Switzerland). The muscle incision was sutured and the skin wound was closed by surgical glue and wound clips. The animal was then removed from the heating pad and monitored until recovery. To reduce dam rejection for the $P5^{Tx}$ model, mouse pups were dabbed in parents' urine before returning to the cage. Animal survival curves and their median survival were determined by the Kaplan–Meier method in GraphPad Prism 7.
2. **Recombinant human CXCL1 protein treatment**
   Treatment of the $P60^{Tx}$ mice with recombinant Human CXCL1 (rhCXCL1) protein (R&D, 275-GR) was started five days after orthotopic transplantation. Mice were treated with rhCXCL1 protein or PBS at 1 ug/100ul/mice via tail veil injection for eight weeks, once a week, five mice/group.
3. **HypoxyProbe**
   Prepare hypoxyprobe solution(116mg/ml) in 0.9% saline. Inject hypoxyprobe solution intraperitoneally into mice at 60mg/kg. After 1.5 to 2 h injection, harvest tissue for fixation and processing for IHC. Anti-pimonidazole mouse IgG1 monoclonal antibody(MAb1) was used for 1:50 dilution.

**Animal tissue staining**

1. **Animal tissue processing**
   Liver and tumor tissues were fixed in neutral buffered formalin for 1 day at room temperature and submitted to HistoWiz Inc. (Brooklyn, NY, USA) for tissue processing and embedding. Formalin-fixed paraffin-embedded (FFPE) tissue sections were cut at 4 µm and analyzed for direct fluorescence microscopy, H&E staining, IHC, and RNAscope.
2. **IHC**
   IHC was performed based on the standard protocol. Antibodies used included anti-vimentin (Abcam, ab92547, 1:1000); anti-αSMA (Abcam, ab124964, 1:1000); and anti-desmin (Abcam, ab32362, 1:2000).
3. **RNAscope**
   RNAscope in situ hybridization of *Cxcl1*, *Hif1α* and *αSMA* mRNA transcripts was performed according to the manufacturer's protocol (Advanced Cell Diagnostics, Hayward, CA, USA).

**Cell lines and culture**

1. **Human HB cell lines**

Human hepatoblastoma cell line HepG2 was purchased from the American Type Culture Collection (ATCC) and cultured in Dulbecco's modified Eagle's medium (DMEM) containing 1% penicillin-streptomycin (P/S), 1% L-Glutamine, and 10% FBS. Human HB214 cell line was acquired from XenTech (Paris, France) and cultured in Advanced DMEM/F12 containing 1% P/S, 1% L-Glutamine, 8% FBS, and 20 μM Rock inhibitor.

2. **Human HSC cell lines**
The human immortalized HSC line LX2 was purchased from Millipore Sigma (Cat. No SCC064. St Louis, Missouri, USA) and cultured in DMEM containing 1% P/S, 1% L-Glutamine, and 2% FBS. Primary human HSCs (HHSteC) were purchased (Cat No. M5300. ScienCell Research Laboratories, San Diego, CA, USA) and cultured in Stellate Cell Medium (Cat No. 5301. ScienCell) supplemented with 1% stellate cell growth supplement (Cat No. 5352. ScienCell), 1% P/S, and 2% FBS.

3. **Normoxia and hypoxia cell culture**
When not specified, cells were cultured under the normoxia condition with 21% $O_2$ and 5% $CO_2$ at 37°C. For hypoxia culture, cells were maintained under 1% $O_2$ generated by flushing a 94% N2/5% $CO_2$ mixture into a Heracell VIOS 160i $CO_2$incubator with Cell Locker System (Thermo Fisher Scientific, Waltham, MA, USA).

##### Generation of CXCR2 KO cell lines by the CRISPR/Cas9 system

*CXCR2*$^{KO}$ HepG2 and HB214 cells were generated via a CRISPR/Cas9 approach. The sgRNA sequence 5′-AAAAATGGAAGATTTTAACANGG-3′ is located at the start codon of *CXCR2* gene.

##### Immunoblotting

Whole-cell lysates were extracted in RIPA plus 100× inhibitor cocktail and 100× EDTA. Twenty μg of protein was loaded to a NuPAGE™ 4 to 12% Bis-Tris, 1.0, Protein gel (Invitrogen, Waltham, MA) for immunoblotting. Antibodies used included anti-CXCR2 (Abcam, ab217314, 1:500), anti-αSMA (Abcam, ab124964, 1:1000), anti-vimentin (Abcam, ab92547,1:1000), anti-desmin (ab32362, 1:1000), anti-Hif1α (Cayman Chemistry, 10006421-1, 1:1000) and anti-vinculin (Cell Signaling Technology , 13901S, 1:2000).

##### Cytokine array assay

Cells were plated onto 10cm dish at a density of 2 x $10^6$ cells/ dish. After 48 hours, the conditioned media from different cell lines were collected and centrifuge to remove any dead or floating cells. Proinflammatory factor levels were measured using human Cytokine Array C5 (RayBiotech, GA, USA) according to the manufacturer's instructions. The intensities of signals were quantified, normalized to positive control. The culture supernatants were also applied onto Human CXCL1/GRO alpha Quantikine ELISA Kit from R&D, following the manufacturer's protocol.

##### Cellular fucntional assays

1. **Haptotaxis Assay**
Cell migration ability was determined by CytoSelect™ 24-Well Cell Haptotaxis Assay (8 μm, Collagen I-Coated, Colorimetric Format; Cell Biolabs, Inc). The chemoattractant was CXCL1 secreted by LX-2 cells. About $1\times10^5$ prepared tumor cells were added into the chamber and $2.4\times10^5$ LX-2 cells were seeded in the well, incubated for 24 h at 37ºC. The cells on the upper surface were wiped with cotton swabs, while the cells that migrated through the filter pores were stained with cell stain solution for 10 min. The stained cells were observed and photographed under an inverted microscope. Cells from five randomly selected fields were counted using Image J.

2. **Apoptosis assay**
   Cells were seeded in 24-well plates with 5 x $10^4$ in each well. After the treatment, cells were incubated in the cultured medium with 2 μM NucView 530 Caspase-3 Substrate(Biotium, Fremont, CA). After 30 minutes, cells were imaged by fluorescence microscopy. Cells from five randomly selected fields were counted using Image J.

3. **Two-chamber migration assay**
   A two-chamber culture insert was placed in the μ-slide 8-well Grid-500 culture slides (ibidi, Gräfelfing, Germany) and the culture surface was precoated with 1% GFR matrigel at 37°C for 24 hr. Tumor and HSC cells were seeded in various combinations, as indicated in the figures. The total number of cells in each chamber was kept at $4.25\times10^4$ cells, and the mixing ratio of T: LX-2 was 4:1. LX-2 were labeled with CellTracker$^{TM}$ Red CMTPX (Thermo Fisher). The cocultures were maintained for 14 days, and the culture medium was refreshed every 3–4 days. The culture medium was a mixture of 80% tumor medium and 20% Stellate Cell Medium (ScienCell). Every 7 days scanned the plates with Lionheart machine (Biotech). Cells from five randomly selected fields were counted using Image J.

All the experiments were repeated >= three time.

##### Quantitative real-time PCR

Total RNA was isolated by a Qiagen kit according to the manufacturer's instructions. A total of 2 μg RNA was subjected to reverse transcription to synthesize cDNA using SuperScript III First-Strand Synthesis SuperMix for qRT-PCR (Invitrogen,11752250). Quantitative real-time PCR for CXCL1 and CXCR2 together with S18 was performed with SYBR Green PCR Master Mixture Reagents (Applied Biosystems) on the ABI PRISM 7900 system (Applied Biosystems). Paired primers were used as follows (5′ to 3′): hCXCR2, forward: CCTGTCTTACTTTTCCGAAGGAC; reverse: TTGCTGTATTGTTGCCCATGT; hCXCL1, forward: AACCGAAGTCATAGCCACAC; reverse: GTCACTGTTCAGCATCTTTTCG; S18, forward: TGTGCCGCTAGAGGTGAAATT ;reverse: TGGCAAATGCTTTCGCTTT .

##### Total mRNA sequencing and analysis

Total RNA library was constructed using Illumina TrueSeq stranded mRNA library prep kit and sequenced using the HiSeq 2000/2500 or NovaSeq 6000 platform (2 x 101- bp pair-end reads). On average, we achieved at least 20x coverage for more than 30% of the transcriptome. Gene expression was quantified by STAR[1] (ver. 2.6.0b) under default parameters with the human genome (GRCh37) and annotation file (Gencode v19).[2, 3] Differential expression was performed by ABSSeq under aFold module.[4] Geneset enrichment analysis was performed by GSEA.[5]

##### Single-cell RNA-sequencing (sc-RNAseq) and analysis

1. **Liver tissue preparation and single cell dissociation**
   One P5$^{Tx}$ mouse and one P60$^{Tx}$ mouse were euthanized 42 days after intrahepatic injection. The tumor and surrounding liver tissue were removed and dissociated with papain digestion solution. Briefly, the tumor were minced and then enzymatically digested in advanced DMEM/ F-12 (Thermo Fisher Scientific, Waltham, MA), containing 10 U/mL Papain (Worthington, Lakewood, NJ), 1 mmol/L N-acetyl cysteine (Sigma-Aldrich, St. Louis, MO), 12mg/mL DNase I (Sigma-Aldrich). The mixture was incubated at 37°C for about 30 min for enzymatic digestion. The additional mechanical digestion was performed by gently pipetting up and down. The cell suspensions were then passed through a 100-μm strainer and washed twice with 5ml of cold Advance DMEM/F12 and centrifugated at 300g for 5min at 4°C.

2. **Droplet-based single-cell RNA sequencing**

The single cells dissociated from the liver and tumor tissues were processed by the Chromium Single Cell Platform using the Chromium Single Cell 3' Library and Gel Bead Kit v2 (10X Genomics) as per the manufacture's protocol. Briefly, the single cells were washed once with PBS plus 0.04% BSA and counted with = a Luna-FL Fluorescence Cell Counter (VitaScientific). Then the cells were added to each lane of the chip and partitioned into Gel Beads in emulsion in the Chromium Controller. The cells were lysed, barcoded, and reversely transcribed, followed by amplification, fragmentation, adaptor attachment and indexing. The libraries were sequenced by Novaseq 6000 (Illumina).

3. **Pre-processing of scRNA-seq data**
To distinguish the human cancer cells from the mouse TME cells, the sequencing data of each sample were aligned to GRCh37(hg19) and GRCm38(mm10) reference Genomes separately using the Cell Ranger Single-Cell Software Suite (v2.0.1, 10X Genomics), and the unique molecular identifiers (UMIs) were estimated with the default parameters. The UMI files of the 4 samples ($P5^{Tx}$ Liver, $P5^{Tx}$ Tumor, $P60^{Tx}$ Liver and $P60^{Tx}$ Tumor) were merged for both human cancer cells and mouse liver cells with an in-house script. The Seurat (V-3.2.2)[6] R package was then introduced to remove the low-quality cells following these criteria: human cancer cells expressed < 1000 or > 8000 genes or mitochondrial gene content > 15% of total UMIs; mouse liver cells expressed <200 or >3000 genes or mitochondrial gene content > 10% of total UMIs.

4. **Dimensionality reduction and clustering of scRNA-seq data**
For data normalization, the UMI count of each gene was divided by the total UMI counts of the cell and scaled the library size to 1,000,000, followed by a natural log plus 1 transformation "NormalizeData"). Then the normalized data was further scaled so that the mean expression across cells is 0 and the variance is 1 ("ScaleData"). Principle component analysis (PCA) was employed for dimensionality reduction ("RunPCA"). To determine the number of components to choose from, a resampling test inspired by the JackStraw procedure ("JackStraw" and "ScoreJackStraw") was implemented and the first 15 principal components were picked, which explained sufficient observed variance. To identify the clusters from the reduced dimensional space, a k-nearest neighborhood (KNN) graph was constructed based on the Euclidean distance in PCA space and refined the edge weights among cells based on the shared overlap in local neighborhoods ("FindNeighbors"). The modularity optimization technique named Louvain algorithm was then applied to iteratively group cells together with a resolution parameter of 0.5 ("FindClusters"). Finally, the Uniform Manifold Approximation and Projection[7] was used to visualize the clustering results ("DimPlot").

5. **Cell type annotation**
To annotate the cell type of each cluster, we firstly curated a list of known marker genes for cell lineages in the liver (**Suppl Table S1**) from literature.[8-10] We then calculated the marker score and cell fraction of each cell linage across all clusters using our in-house software package scMINER (v-0.1.0, "draw.marker.bbq"). The marker score is defined as the geometric mean of the expression of the associated marker genes in that cell, and the cell fraction is defined as the proportion of cells of which the geometric mean of marker gene expression is greater than a given threshold (0.5 by default). The expression of exemplar markers was visualized in UMAP plots by ggplot2 (v-3.3.5)[11] R package to confirm the cell type annotation results.

6. **Heatmap of marker gene signatures of cell lineages**
The normalized and scaled UMI counts by Seurat were used to indicate the expression of genes across cells. The expression matrix of curated marker genes was extracted by an in-house R script. The heatmap was constructed by pheatmap (v-1.0.12) [12]R package. The column-wise scaling was applied.

7. **Differential expression analysis**

Monocle 2 (v-2.14.0)[13] was utilized to conduct the differential expression analysis of cell subpopulations across sample sources. In brief, the gene by cell matrix of raw UMIs was created by Seurat and then subjected to create the new CellDateSet object by Monocle 2 with the negative binomial data distribution ("newCellDataSet"). The size factors ("estimateSizeFactors") and estimated the dispersion values ("estimateDispersions") were calculated using the default methods. The significance of differential expression was then calculated using the default likelihood ratio test. To calculate the fold change, the raw UMI counts were normalized by the library size and scaled to 1,000,000 by NetBID 2 (v-2.0.3)[14] R package, followed by the log2-transformation of scaled UMI plus 1. The geometric mean of log2-transformed expression values was then calculated and the difference of subtraction of each comparison was estimated. The differentially expressed genes (DEGs) were defined by these cutoffs: P-value < 0.01 and log2 Fold Change > 1 (up-regulated) or < -1 (down-regulated). The volcano plot was constructed by ggplot2 (v-3.3.5) R package to visualize the differential expression analysis.

8. **Gene set analysis**

The HALLMARK gene sets from MSigDB database (v-6.1)[5] were used for the three gene set analysis. First, we used the fisher exact test to identify the gene sets significantly enriched by the DEGs defined from the differential expression analysis. A cutoff of P-value < 0.001 was applied. Secondly, we introduced the gene set enrichment analysis (GSEA) to evaluate the change of hypoxia between neonatal and adult mice. The P-value and normalized enrichment score (NES) were calculated by FGSEA(v-1.12.0) [15] R package, and the GSEA plot was constructed by NetBID2 (v-2.0.3, "draw.GSEA"). And thirdly, we employed a systems biology method to measure the hypoxia activity in both neonatal and adult mice. The hypoxia activity was defined as the geometric mean of expression of genes involved in the HALLMARK HYPOXIA gene set. The activity was calculated by NetBID2 (v-2.0.3, “cal.Activity.GS”).

**Retrospective examination of archival HB patient samples**

A search was performed of the pathology archives for cases that met the following criteria: (1) tumor diagnosis of HB of any histologic subtype; (2) tumor/adjacent liver interface of at least 5 mm, as evaluated by examination of an adjacent H&E-stained slide; and (3) available unstained slides or FFPE tissue for staining. Each case of HB metastatic at presentation was paired with a case of non-metastatic HB matched with respect to patient age at diagnosis, presence or absence of pre-resection chemotherapy, and histologic subtype of HB when possible. Available clinical history was recorded. Immunohistochemical staining for αSMA (clone 1A4, Dako) was performed using a clinically validated protocol. All the HB patient samples were obtained under a protocol approved by the Institutional Review Boards at St. Jude Children's Research Hospital.

**Patient cohorts collection and survival analysis**

1. **Pediatric patient cohort from GEO/EMBL-EBI**

We collected the pediatric cancer patient information from the QIAGEN Oncoland (Pediatrics, 2018Q1 release) and identified 6 pediatric cancer types with >20 patients with both gene expression profiles and overall survival data: hepatoblastoma (HB) patient cohort (n=55) from GSE75271[16], ependymoma (EPN) patient cohort (n=65) from GSE50385[17], primitive neuroectodermal tumor (PNET) patient cohort (n=56) from GSE14295[18], pediatric glioblastoma multiforme (pGBM) patient cohort (n=48) from GSE32374[19] and GSE34824[20], medulloblastoma (MB) patient cohort (n=92) from GSE30074[21] and GSE68956[22] and malignant glioma patient cohort (n=80) from EMBL-EBI under the project of E-TABM-1107.[23] The quantile normalization, coupled with log2 transformation, were conducted to generate the gene expression profile of each sample at the probe level. Then the probe-level expression values were further aggreged into gene-level expression values by taking the mean when multiple probes were found for one single gene. The overall survival data were directly captured from the QIAGEN Oncoland. The risk-group information of HB cohort was collected from the supplementary data of the paper.[24]

2. **Pediatric patient cohort from TARGET**
   The TARGET RNA-seq data of 7 childhood cancer types at the gene level were extracted from the QIAGEN Oncoland (LandVersion: TARGET_B38_20170310_v1). The samples with low mapping rates (<70%) were excluded, making 52776 genes across 1319 samples in the final cohort. The log2-transformed FPKM values were used as the gene expression profiles. The Rhabdoid Tumor (RT) samples were excluded in subsequent analysis for lacking survival information.
3. **Adult primary cancer patient cohort from TCGA**
   The TCGA RNA-seq data of 32 human primary cancer types at the gene level were extracted from the QIAGEN Oncoland (LandVersion: TCGA_B38_20190215_v8). The samples with low mapping rates (<70%) were excluded, making 55585 genes across 11125 samples in the final cohort. The log2-transformed FPKM values were used as the gene expression profiles.
4. **Survival analysis**
   Only the samples with both gene expression data and survival information were involved in this survival analysis. Kaplan-Meier estimates were constructed by survival R package (v-3.2-13, "survfit")[25] and visualized by survminer R package (v-0.4.9, "ggsurvplot").[26] The two groups for comparison were defined as the top ("High") and bottom ("Low") quarter of patients with regard to the expression of HIF1A. The p value of two-group comparison was calculated by log-rank test[27]. The association between overall survival and hypoxia activity was constructed using the Cox proportional hazards regression model.[28] The hazard ratio, 95% CI and p values were calculated by survival R package (v-3.2-13, "coxph")[25]. The linerange plots were prepared by ggplot2 (v-3.3.5).[29]

**Statistical analysis**

A two-tailed Student *t* test was performed in GraphPad to compare two independent pairs of groups. A one-way ANOVA was performed when two or more groups were compared. A $P \leq 0.05$ was considered statistically significant.

**Data and code availability**

The raw scRNA-seq data have been deposited in the Gene Expression Omnibus (GEO, Accession number: GSE186027). The codes of the data analysis in this paper are freely available at https://github.com/jyyulab/Heptoblastoma_Cxcl1. The total RNA-seq data have been deposited in GEO (Accession number: GSE186335).

**Supplemental Table S1. List of marker genes for cell lineages in the mouse liver.**

| Cell Type | Marker | Weight |
|---|---|---|
| Endothelial cell | Clec14a | 1 |
| Endothelial cell | Pecam1 | 1 |
| Endothelial cell | Clec4m | 1 |
| Endothelial cell | Icam2 | 1 |
| Endothelial cell | Kdr | 1 |
| Endothelial cell | Egfl7 | 1 |
| Endothelial cell | Hspg2 | 1 |
| Endothelial cell | Emcn | 1 |
| Endothelial cell | Oit3 | 1 |
| Endothelial cell | Fcn3 | 1 |
| Endothelial cell | Clec4g | 1 |
| Erythrocyte | Hba-a1 | 1 |
| Erythrocyte | Hba-a2 | 1 |
| Erythrocyte | Hbb-bs | 1 |
| Erythrocyte | Hbb-bt | 1 |
| Erythrocyte | Gypa | 1 |
| conventional dendritic cell | Itgax | 1 |
| conventional dendritic cell | Btla | 1 |
| conventional dendritic cell | Cd24a | 1 |
| conventional dendritic cell | Xcr1 | 1 |
| conventional dendritic cell | Irf8 | 1 |
| plasmacytoid dendritic cell | Clec4b1 | 1 |
| plasmacytoid dendritic cell | Pira2 | 1 |
| plasmacytoid dendritic cell | Trem2 | 1 |
| plasmacytoid dendritic cell | Ly6c1 | 1 |
| Macrophage | Adgre1 | 1 |
| Macrophage | Cd163 | 1 |
| Macrophage | Cd68 | 1 |
| Macrophage | Clec4f | 1 |
| Macrophage | Cxcl13 | 1 |
| Macrophage | Cd5l | 1 |
| Macrophage | Fabp7 | 1 |
| Hepatic stellate cell | Acta2 | 1 |
| Hepatic stellate cell | Col1a1 | 1 |
| Hepatic stellate cell | Col1a2 | 1 |
| Hepatic stellate cell | Tagln | 1 |
| Hepatic stellate cell | Tpm2 | 1 |
| Hepatic stellate cell | Colec11 | 1 |
| Hepatic stellate cell | Rgs5 | 1 |
| Hepatic stellate cell | Ecm1 | 1 |
| Epithelial cell | Epcam | 1 |
| Epithelial cell | Alb | 1 |
| Epithelial cell | Ttr | 1 |
| Epithelial cell | Krt18 | 1 |
| Epithelial cell | Krt19 | 1 |
| Epithelial cell | Sox9 | 1 |

### Suppl Figure S1

A

HB214 (HB-PDX derived cell line)

| Host age | Cell # /mouse | Engraftment Rate |
|---|---|---|
| P5 | $5\times10^4$ | 0/3 |
| P5 | $1\times10^5$ | 0/3 |
| P21 | $1\times10^6$ | **5/5** |
| P60 | $1\times10^6$ | 0/5 |

B

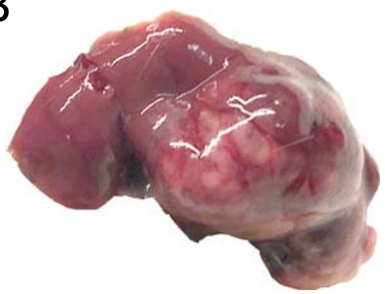

C

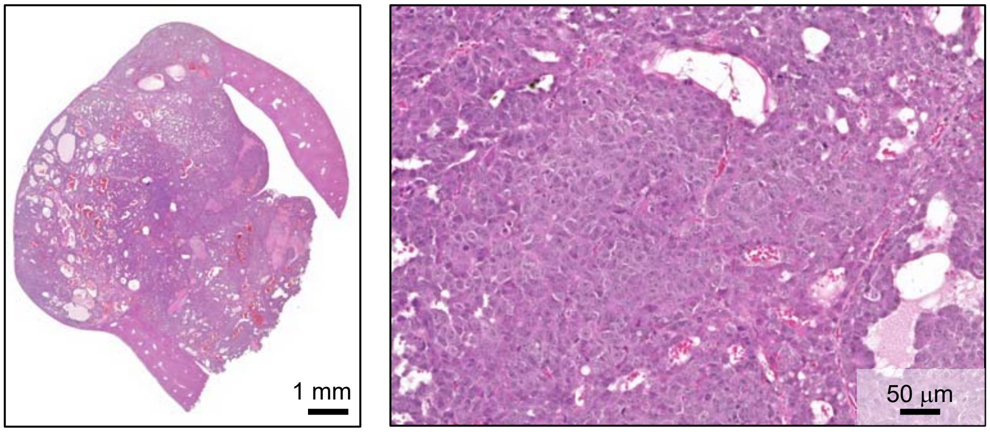


**Suppl Figure S1. HB214 cells are more tumorigenic in the immature liver than in the adult liver.**

**(A)** A table showing the age of the host, number of cells used, and tumor engraftment rate of the HB214-transplanted NSG mice.

**(B)** Gross image of a liver tumor from a mouse transplanted with HB214 at P21.

**(C)** Low- and high-magnification H&E images of a $P21^{Tx}$ HB214 tumor.

### Suppl Figure S2

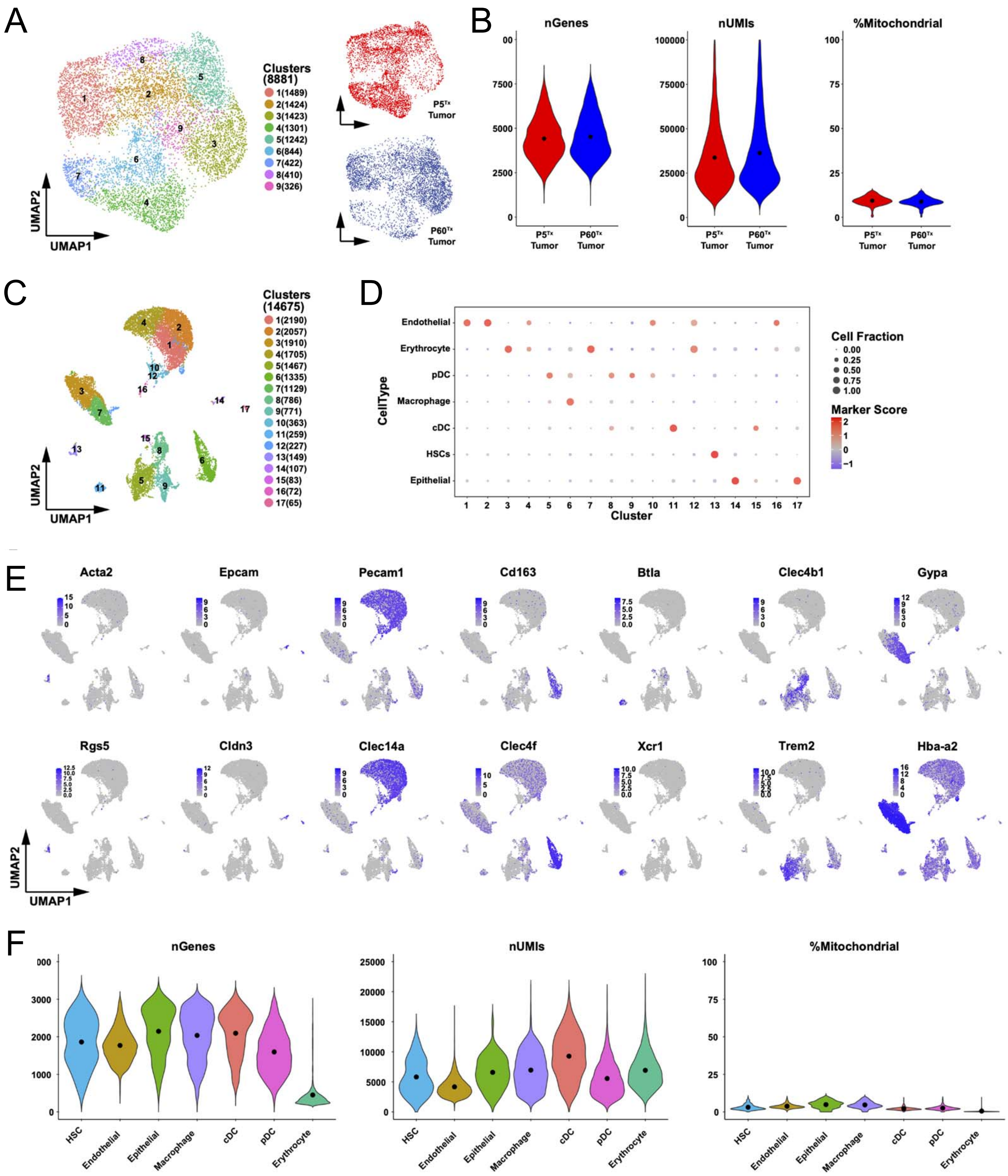

**Suppl Figure S2. Quality control and cell type annotation of sc-RNAseq data.**

**(A)** Clustering of 8881 bulk tumor cells from the HepG2 $P5^{Tx}$ (red) and $P60^{Tx}$ (blue) model combined.

**(B)** Violin plots showing the number of genes (left), the number of UMIs (middle) and the percentage of mitochondrial genes (right) across the bulk tumor cells from the $P5^{Tx}$ and $P60^{Tx}$ model. Black dots in the violin plots denote the mean values.

**(C)** Clustering of 14675 tumor microenvironment cells from the $P5^{Tx}$ and $P60^{Tx}$ model combined.

**(D)** Dot plot showing the cell type annotation results of tumor microenvironment cells. The circle size indicates the fraction of cells of which the mean expression of marker genes greater than the default cutoff. The color indicates the marker score of each cell type across all clusters.

**(E)** Violin plots showing the number of genes (left), the number of UMI counts (middle) and the percentage of mitochondrial genes (right) across seven cell types identified from the tumor microenvironment cells of the $P5^{Tx}$ and $P60^{Tx}$ model. Black dots in the violin plots denote the mean values.

**(F)** UMAP plots showing the expression of exemplar marker genes for each cell type.

### Suppl Figure S3

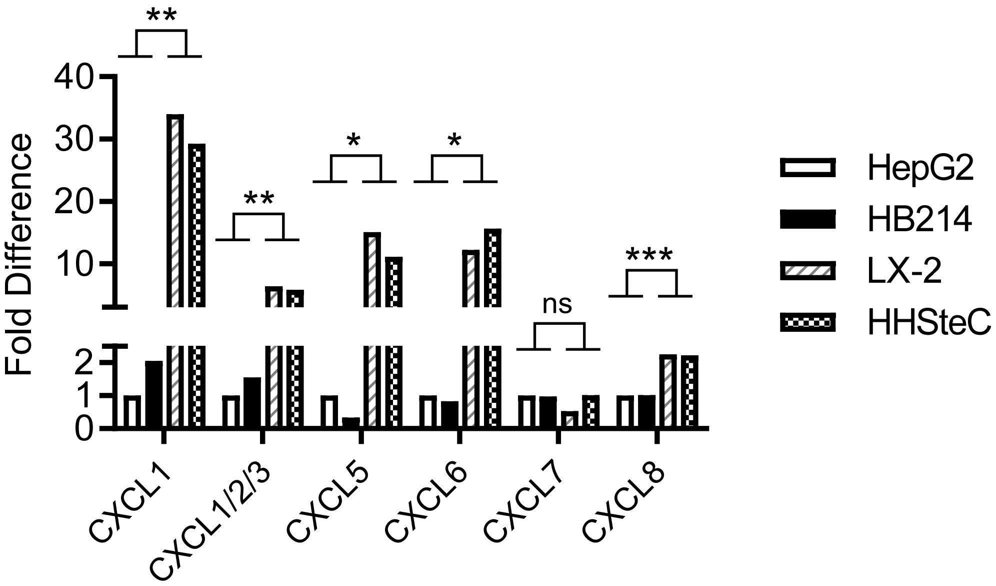


**Suppl Figure S3. Multiple CXCR2 ligands are secreted at a higher level in aHSCs than HB tumor cells in vitro.**

Image-based quantification and fold comparison of the indicated cytokine signals shown in **Figure 4A.** *P* values were calculated by unpaired two-tailed *t*-test: ns, not significant, *<0.05; ** <0.01; *** <0.001.

### Suppl Figure S4

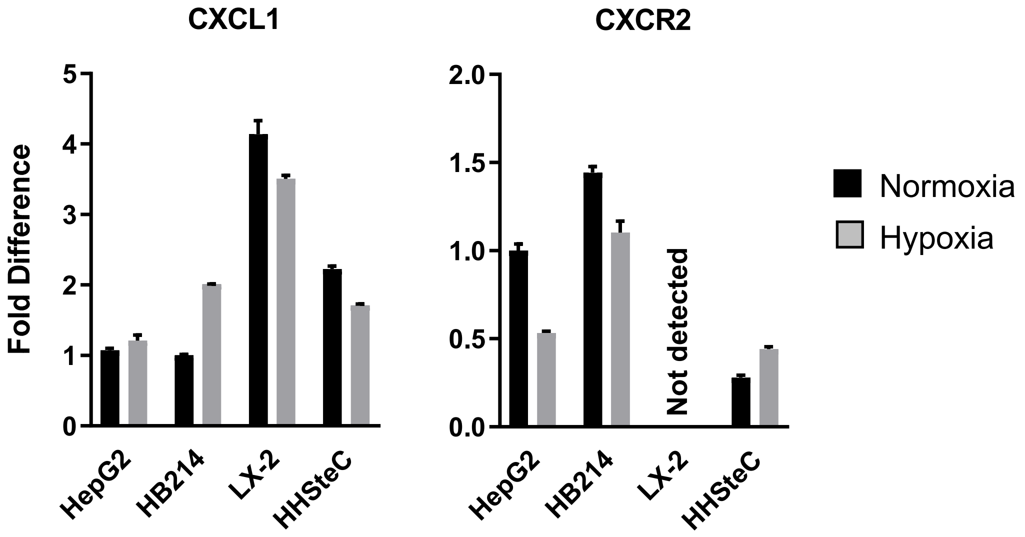


**Suppl Figure S4. *CXCl1* and *CXCR2* expression in HSC and HB cells cultured under normoxia and hypoxia in vitro.**

Real-time quantitative PCR of *CXCl1* and *CXCR2* in the HSC and HB tumor cells cultured under normoxia and hypoxia. No statistical analysis was performed due to the inconsistent changes induced by hypoxia among the cell lines.

### Suppl Figure S5

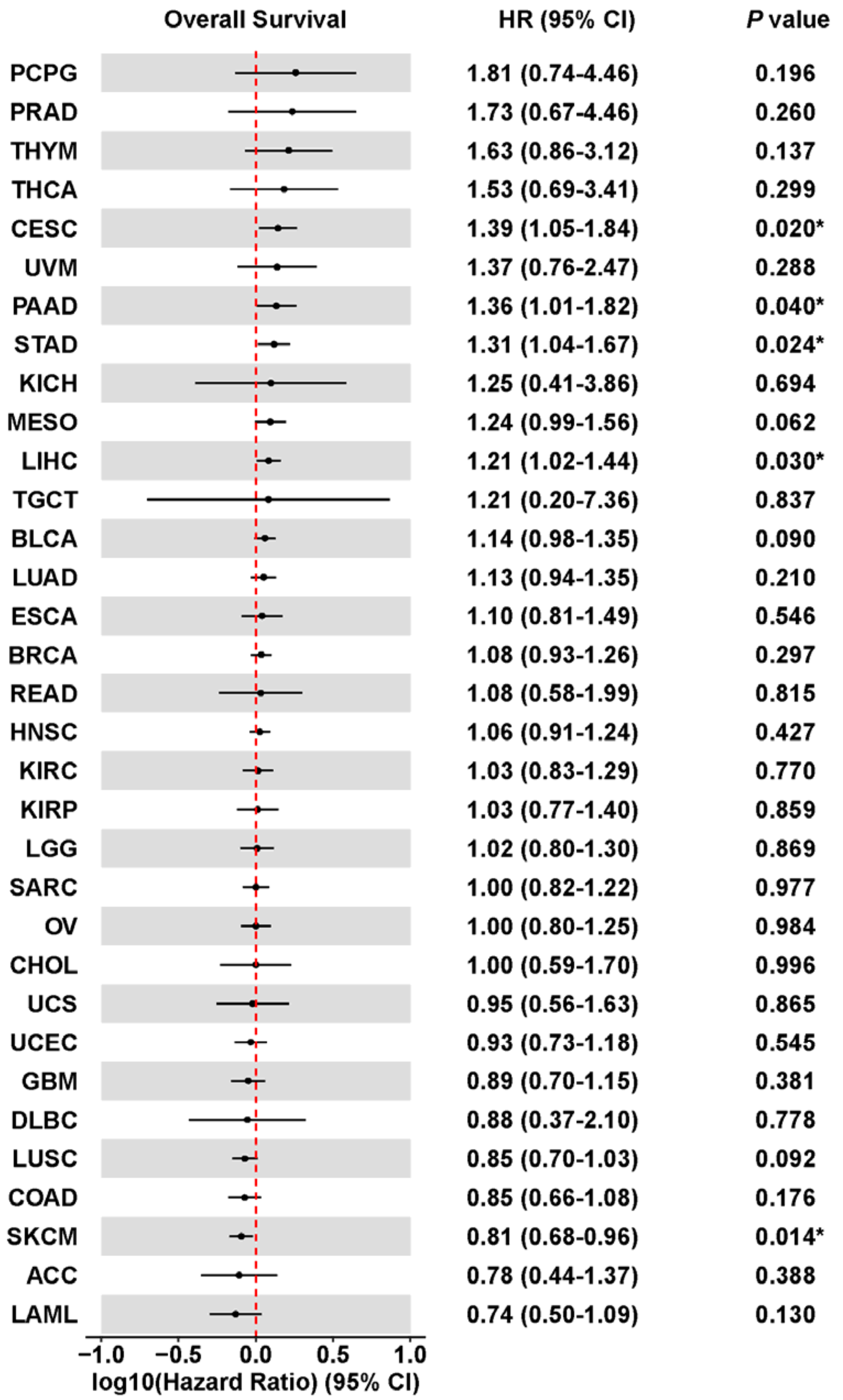

**Suppl Figure S5. Association between *HIF1A* expression and the overall survival of 33 types of adult primary cancer patients from TCGA cohort.**

ACC: Adrenocortical carcinoma; BLCA: Bladder Urothelial Carcinoma; BRCA: Breast invasive carcinoma; CESC: Cervical squamous cell carcinoma and endocervical adenocarcinoma; CHOL: Cholangiocarcinoma; COAD: Colon adenocarcinoma; DLBC: Lymphoid Neoplasm Diffuse Large B-cell Lymphoma; ESCA: Esophageal carcinoma; GBM: Glioblastoma multiforme; HNSC: Head and Neck squamous cell carcinoma; KICH: Kidney Chromophobe; KIRC: Kidney renal clear cell carcinoma; KIRP: Kidney renal papillary cell carcinoma; LAML: Acute Myeloid Leukemia; LGG: Brain Lower Grade Glioma; LIHC: Liver hepatocellular carcinoma; LUAD: Lung adenocarcinoma; LUSC: Lung squamous cell carcinoma; MESO: Mesothelioma; OV: Ovarian serous cystadenocarcinoma; PAAD: Pancreatic adenocarcinoma; PCPG: Pheochromocytoma and Paraganglioma; PRAD: Prostate adenocarcinoma; READ: Rectum adenocarcinoma; SARC: Sarcoma; SKCM: Skin Cutaneous Melanoma; STAD: Stomach adenocarcinoma; TGCT: Testicular Germ Cell Tumors; THCA: Thyroid carcinoma; THYM: Thymoma; UCEC: Uterine Corpus Endometrial Carcinoma; UCS: Uterine Carcinosarcoma; UVM: Uveal Melanoma.

HR, 95% CI and *P* values were calculated by univariate Cox proportional-hazards model. *P* values * < 0.05, ** < 0.01, and *** < 0.001.
